# Formation of chromosomal rearrangements in *Saccharomyces cerevisiae* diploids through regionally-biased non-allelic homologous recombination

**DOI:** 10.1101/2025.05.08.650247

**Authors:** Sean A. Merriman, Mary J. Chapman, Joseph A. Stewart, Camryn D. Schmelzer, Rabab S. Sharif, Megan J. Hemmerlein, Christopher M. Puccia, Guilherme M. de Mattos, Mackenzie A. Wienke, Deborah A. Cornélio, Matthew Dilsaver, Ruth A. Watson, Juan Lucas Argueso

## Abstract

In earlier studies, we optimized an assay system for the genome-wide detection of copy number variation (CNV) in diploid *Saccharomyces cerevisiae* cells, based on selection for formaldehyde plus copper (FA+Cu) resistance conferred by the amplification of a dosage-dependent reporter cassette, *SFA1-CUP1*. Our analyses identified a robust bias for terminal deletions of the right arm of Chr7 (Chr7R) associated with unbalanced translocations. This bias was observed at approximately constant strength across all three sites where the amplification reporter cassette was inserted, in CNV-carrying yeast clones derived both spontaneously and from mutagen-induced recombinogenic conditions. We conducted allelic mitotic recombination experiments to investigate the possibility of the presence of a fragile site on Chr7R, but the results disfavored this model, and instead indicated that the Chr7R bias applies only to non-allelic rearrangements. We validated the existence of a CNV formation bias at Chr7R through an orthologous NAHR competition approach that was independent of selection for FA+Cu resistance. Finally, we showed the in contrast to its high participation in NAHR as a recipient sequence, Chr7R becomes amplified as a translocation donor less frequently than other comparable regions of the genome. To begin unraveling the cause of this unusual behavior, we evaluated the effect of a set of candidate genes involved in chromatin mobility and sister chromatid cohesion on the rearrangement spectra involving Chr7R. We found that deletion mutations in some of these genes, particularly *SAP30*, attenuated the biased NAHR behavior. Taken together, our results suggested that although Chr7R is not inherently more prone to DNA breakage than other regions, once a DNA lesion is formed there, it has a higher propensity to undergo inappropriate repair leading to a chromosomal rearrangement.

## INTRODUCTION

In the last decades, the field of genomic medicine has experienced remarkable growth made possible by improvements in the throughput and accuracy of DNA sequencing technologies. One of the main breakthroughs that followed was the discovery that many of the genetic differences that exist between individuals are variations in the number of copies of their genes (Zhang et al. 2009). Such gene copy number variations (CNVs) are a particularly important component of the altered genomes of cancer cells (Beroukhim et al. 2010; Steele et al. 2022). Despite the importance of CNVs to genetic variation, genomic disorders and cancer development, our understanding of the fundamental mechanisms that trigger these large-scale mutations is still incomplete (Mani and Chinnaiyan 2010). Genetic assays needed to detect and experimentally characterize their formation remain limited in mammalian cells. Much of the foundational knowledge in this area has, and continues to be established in simpler eukaryotic model organisms, such as the budding yeast *Saccharomyces cerevisiae*, in which powerful cell-based recombination assay systems can be leveraged (Klein et al. 2019), and the detailed analysis of the resulting chromosomal rearrangement products can be achieved relatively practically and affordably (Heasley et al. 2021).

Structural rearrangements of the genome are initiated by DNA double-strand break (DSB) lesions that can arise from both endogenous and exogenous sources (Arlt et al. 2012). Studies in *S. cerevisiae* have provided valuable insights into the homologous repair pathways used to process DSB lesions and the repair and mis-repair outcomes that result from these pathways (Symington et al. 2014; Putnam and Kolodner 2017). The canonical two-ended Homologous Recombination Double Strand Break Repair pathway (HR-DSBR) is recognized as the most conservative pathway, especially when using an intact, allelic sequence template to repair a damaged chromosome. Use of the allelic template present in the sister chromatid leads to accurate and fully restorative repair, whereas use of the allelic template present in the homolog preserves the original chromosomal structure, but can lead to copy neutral loss of heterozygosity (cnLOH) (Dutta and Schacherer 2025).

While the primary HR-DSB repair pathway above leads to conservation of the chromosome structure and DNA copy number, some of its homology-mediated variations can lead to mis-repair associated with gains or losses of large genomic segments, specifically when a non-allelic repeat sequence is used as the template (Guirouilh-Barbat et al. 2014; Savocco and Piazza 2021). For example, if a DSB occurs between two direct repeats present in close proximity, resection of both ends can lead to annealing of exposed single-strands in the Single Strand Annealing (SSA) mechanism (Paques and Haber 1999), resulting in interstitial deletion of the region between the repeats. If one of the two ends of the DSB is lost, Break-Induced Replication (BIR) may be used to salvage the remaining end (Kockler et al. 2021), but if the ensuing homology search engages a non-allelic template, the mis-repair outcome will include both amplification and deletion of genomic segments. Finally, the canonical two-ended HR-DSB repair pathway can also lead to chromosomal rearrangements if a homologous non-allelic template is used to repair a DSB. In this case, if the recombination intermediate is resolved in the crossover orientation leading to exchange of the regions flanking the repeats, then a wide range of structural variation types can be formed (Argueso et al. 2008; Reitz et al. 2023), including both reciprocal and non-reciprocal translocations associated with amplifications and deletions.

Among the various triggers to the formation of structural genomic rearrangements (Mani and Chinnaiyan 2010), the spatial organization of the genome has emerged as an important determinant of DSB repair in organisms ranging from yeast to humans. It has been shown that cancer-causing translocations in human cells are correlated with spatial proximity of the loci which recombine to create them (Roix et al. 2003; Roukos and Misteli 2014). The territories chromosomes occupy in the yeast nucleus have been established as a limit on which sequences recombine (Taddei and Gasser 2012; Agmon et al. 2013; Miné-Hattab and Rothstein 2013). Chromosomes in the *S. cerevisiae* genome are known to organize in the “Rabl-like” configuration, in which the centromeres of all chromosomes are clustered together at one nuclear pole while the telomeres cluster together at the opposite pole of the nucleus (Duan et al. 2010). This arrangement results in loci of similar distance from the centromere or telomere occupying similar regions, and recombining more frequently as a result (Khrameeva et al. 2016; Haber 2018).

While spatial organization of the genome places constraints on which loci recombine, the nucleus is not static (Garcia Fernandez and Fabre 2022; Chiolo et al. 2025). In yeast, global chromatin mobility increases in response to DSB formation, and the loci of a DSB itself undergoes the most drastic increase in mobility (Dion et al. 2012; Mine-Hattab and Rothstein 2012; Garcia Fernandez et al. 2022). This mobility increase enhances the efficiency of the homology search in HR. The nearest and most ideal template to repair a DSB in mitotic cells is the sister chromatid, and the close proximity between a damaged locus and a sister chromatid is maintained by sister chromatid cohesion (SCC). Defects in SCC, such as loss of Scc1, a protein which links cohesin rings together, have been reported to decrease sister chromatid recombination and in turn elevate non-allelic recombination (Dion et al. 2013; Mackenroth and Alani 2021). SCC is therefore important for promoting accurate repair of DSBs and avoiding the generation of deleterious chromosomal rearrangements (Piazza et al. 2021; Phipps and Dubrana 2022).

What mechanisms determine if two non-homologous chromosomes are more likely to interact as translocation partners? Do the unique positioning, movement, and mobility of different chromosomes allow certain rearrangements to occur more often than others? What proteins are involved in maintenance of these dynamics? We sought to answer some of these outstanding questions in the context of a striking bias in translocation partners that we have observed in our prior studies (Stanton 2012; Sharif 2014). We have used *S. cerevisiae* as a model in which to study CNV-generating mechanisms, utilizing a *SFA1^V208I^*-*CUP1* amplification reporter cassette, conferring dosage-dependent formaldehyde and copper resistance, respectively (Zhang et al. 2013; Klein et al. 2019). With this system, we identified a specific region of the *S. cerevisiae* genome (the right arm of chromosome 7; Chr7R) that is much more susceptible to deletion during formation of unbalanced, nonreciprocal translocations that amplify other chromosome regions. The junctions of these translocations are often marked by the presence of dispersed repeats, strongly implying a non-allelic homologous recombination (NAHR) mechanism. However, the reason for preference of Chr7R as a translocation recipient remains unclear. In this study, we independently validated the existence of such Chr7R bias for deletion in NAHR translocations, compared its behavior as both translocation donor versus a recipient sequence, and interrogated the participation in this process of a small set of candidate genes. We showed that this unusual behavior is likely the result of a NAHR directionality repair bias, and that it can be attenuated by loss of genes important for maintaining chromatin mobility and sister chromatid cohesion.

## RESULTS

### Initial observation of frequent deletions spanning the right arm of Chr7

We first noted a distinct enrichment in Chr7 right arm (Chr7R) deletions in our earlier work where formaldehyde and copper resistant (FCR) clones were derived from three different experimental parent diploid strains, in which the *SFA1^V208I^*-*CUP1* amplification reporter cassette was inserted at loci on either Chr4R, Chr5R, or Chr15R (Fig. 1) (Stanton 2012; Sharif 2014; Klein et al. 2019). The chromosomal aberrations present in all FCR clones isolated through that work were characterized in detail using pulse field gel electrophoresis (PFGE) and microarray-based comparative genomic hybridization (array-CGH), and in several cases also by nucleotide sequencing analyses at the rearrangement junctions through Nanopore or PCR-Sanger. The predominant rearrangements present in them were unbalanced translocations mediated by non-allelic homologous recombination (NAHR), in which the donor DNA sequence was a repeat present in the amplified chromosomal arm (*i.e.*, the arm with *SFA1^V208I^*-*CUP1* reporter insertion), and the recipient sequence (*i.e.*, the presumed site of the precursor DSB that triggered NAHR repair) was a homologous DNA repeat present at the endpoint of a Chr7R terminal deletion tract.

**Figure 1.**
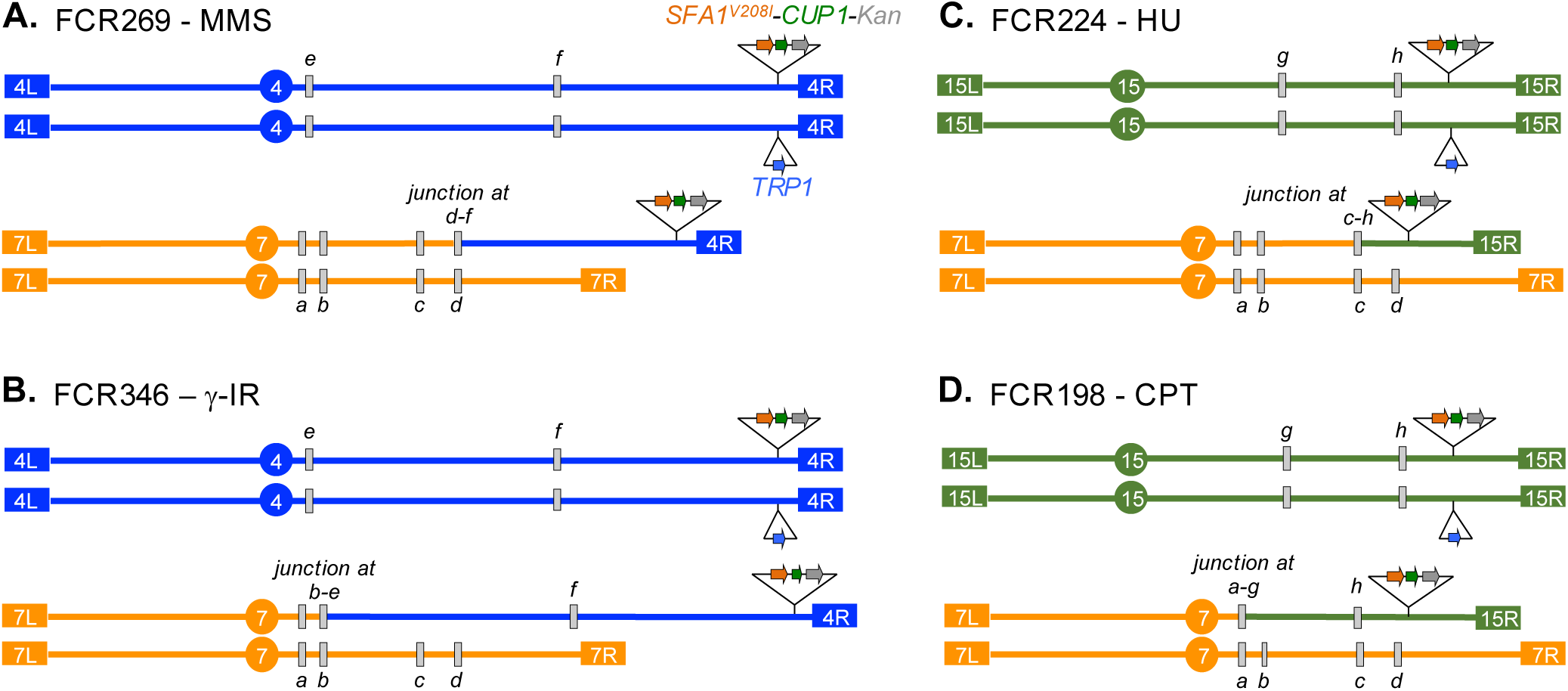
Schematic karyotypes of representative FCR clones carrying NAHR-mediated non-reciprocal translocations amplifying Chr4R or Chr15R and deleting Chr7R. A. Clone FCR269, derived from the experimental diploid JAY654 with the *SFA1^V208I^-CUP1* reporter inserted at Chr4R, isolated after exposure to methyl methanesulfonate (35 mg/ml; MMS). B. FCR346, also derived from the JAY654 Chr4R reporter parent strain, after exposure to γ ionizing radiation (200 Gy; γ-IR). C. FCR224, derived from the experimental diploid JAY685 with the *SFA1^V208I^-CUP1* reporter inserted at Chr15R, isolated after exposure to hydroxyurea (50 mM; HU). D. FCR198, also derived from the JAY685 Chr15R reporter parent strain, after exposure to camptothecin (15 μg/ml; CPT). The gray boxes labelled with lower-case italicized letters (*a* through *f*) on chromosomes indicate the regions containing DNA repeat sequences involved in the NAHR rearrangements, and the respective translocation junctions found at each rearrangement. Chr7R repeat regions: *a*. *YGRWTy1-1*; *b*. *YGRCTy1-2* and *YGRCTy2-1*; *c*. *YGRWsigma6*; *d*. *YGRWTy2-2* and *YGRCTy1-3*. Chr4R repeat regions: *e*. *YDRCTy2-1*; *f*. *YDRWTy2-3* and *YDRCTy1-3*. Chr15R repeat regions: *g*. *YORWTy1-2*; *h*. *YORWsigma3*. Note that there are many additional repeat sites present in these chromosomes which were not involved in the example rearrangements shown. These uninvolved repeat sites are not displayed in the figure for simplicity. The approximate positions of *SFA1^V208I^-CUP1-Kan* CNV reporter (orange-green-gray arrows) insertion sites are shown for Chr4R (between *PLM2* and *SAM2*) and Chr15R (between *RPL20B* and *SSP4*). The *TRP1* auxiliary marker (blue arrow) was inserted at the respective allelic positions on the other Chr4 or Chr15 homologs.

After pooling the FCRs derived from these three parent strains, encompassing clones recovered from normal growth conditions as well as those resulting from exposure to mutagens (HU, MMS, CPT, and γ-IR), we found that roughly half of the FCRs exhibited deletions tracts on Chr7R, while the remaining FCRs had deletions in other chromosome arms (Table 1). Notably, considering the entire set of FCR clones in aggregate, the deletion tract endpoints were not clustered at a discrete site or narrow region on Chr7R. Instead, they were broadly distributed along a large (∼500 Kb) segment of Chr7R, with higher incidence of endpoints at full length Ty element insertions, and additional instances at shorter and lower copy dispersed repeat classes such as solo LTRs and tRNAs. Figure 1 illustrates this diversity of translocation junctions, showing four independent examples of unbalanced translocations, each with a Chr7R deletion endpoint at a different repeat sequence site (gray boxes *a*, *b*, *c*, and *d*). However, a degree of clustering was observed within FCR subsets depending on the chromosome arm where the *SFA1^V208I^-CUP1* amplification reporter cassette was inserted. Specifically, FCRs derived from the Chr4R reporter strain had 36 of 43 deletion endpoints at the region around *YGRCTy1-3* and *YGRWTy2-2* (position ∼818 kb; gray box *d* in Fig. 1), whereas FCRs derived from the Chr15R reporter strain had 14 of 22 deletion endpoints at the region around *YGRCTy1-2* and *YGRCTy2-1* (position ∼573 kb; gray box *b* in Fig. 1).

**Table 1.** Biased regional involvement in non-reciprocal inter-chromosomal translocations: Reporter arm amplifications and associated arm deletions.

| Terminal arm amplifications & CNV reporter insertion sites | Mutagen exposure | <sup>a</sup> Associated terminal arm deletions in FCRs |  |
| --- | --- | --- | --- |
|  |  | Chr7R | All other arms combined |
| Strain JAY654 - Chr4R<br>*SFA1<sup>V208I</sup>-CUP1* inserted between *PLM2* and *SAM2* | uninduced | 11 | 10 |
|  | HU | 6 | 3 |
|  | CPT | 9 | 2 |
|  | MMS | 9 | 3 |
| | γ-IR | 8 | 4 |
| Strain JAY538 - Chr5R<br>*SFA1<sup>V208I</sup>-CUP1* inserted between *DDI1* and *UBP5* | uninduced | 2 | 3 |
|  | HU | 3 | 3 |
|  | CPT | 4 | 4 |
|  | MMS | 2 | 3 |
| | γ-IR | nt | nt |
| Strain JAY685 - Chr15R<br>*SFA1<sup>V208I</sup>-CUP1* inserted between *RPL20B* and *SSP4* | uninduced | 8 | 7 |
|  | HU | 3 | 6 |
|  | CPT | 2 | 5 |
|  | MMS | 3 | 7 |
| | γ-IR | 4 | 12 |
| All reporter sites | All conditions | 74 | 72 |

<sup>a</sup> The numbers indicate the counts of FCR clones obtained from the respective strains and growth conditions, which carried non-reciprocal translocations, as determined by array-CGH, with a terminal deletion of Chr7R or a terminal deletion at any/all other chromosome arms combined (e.g., Chr2R, Chr13R, Chr14L, etc). Combined datasets from Stanton 2012 and Sharif 2014.
nt. not tested; HU. Hydroxyurea 50 mM; CPT. Camptothecin 15 μg/ml; MMS. Methyl methanesulfonate 35 μg/ml; γ-IR. Gamma ionizing radiation combined 50 or 200 Gy.

Because our initial set of genomically-analyzed FCR clones was relatively small and derived from a variety of growth conditions (*i.e.* with and without induced DNA damage), we sought to reproduce and validate the Chr7R deletion bias in a larger set of clones obtained from homogeneous uninduced conditions. To do this more efficiently, we modified the experimental strains to enable a faster phenotypic assessment of the presence of Chr7R deletions among FCR clones, bypassing the need for DNA-based genome-wide analyses. The modified experimental parent strains used in this case possessed the same *SFA1^V208I^-CUP1* amplification reporter cassette as before, inserted on Chr4R or Chr15R (Fig. 2A and 2B, respectively), but in addition, they also contained unique marker genes inserted at allelic telomere-proximal sites on each Chr7R homolog: *URA3* and Hph. With Chr7R marked in this way, following isolation of FCR clones carrying Chr4R or Chr15R amplification events, a concurrent deletion of either Chr7 right arm marker could be inferred through growth phenotypes associated with loss of *URA3* or Hph. While this approach was informative about the presence of Chr7R deletions among FCR clones, it did not offer any details on the discrete deletion endpoints or the classes of the chromosomal aberrations present in them.

**Figure 2.**
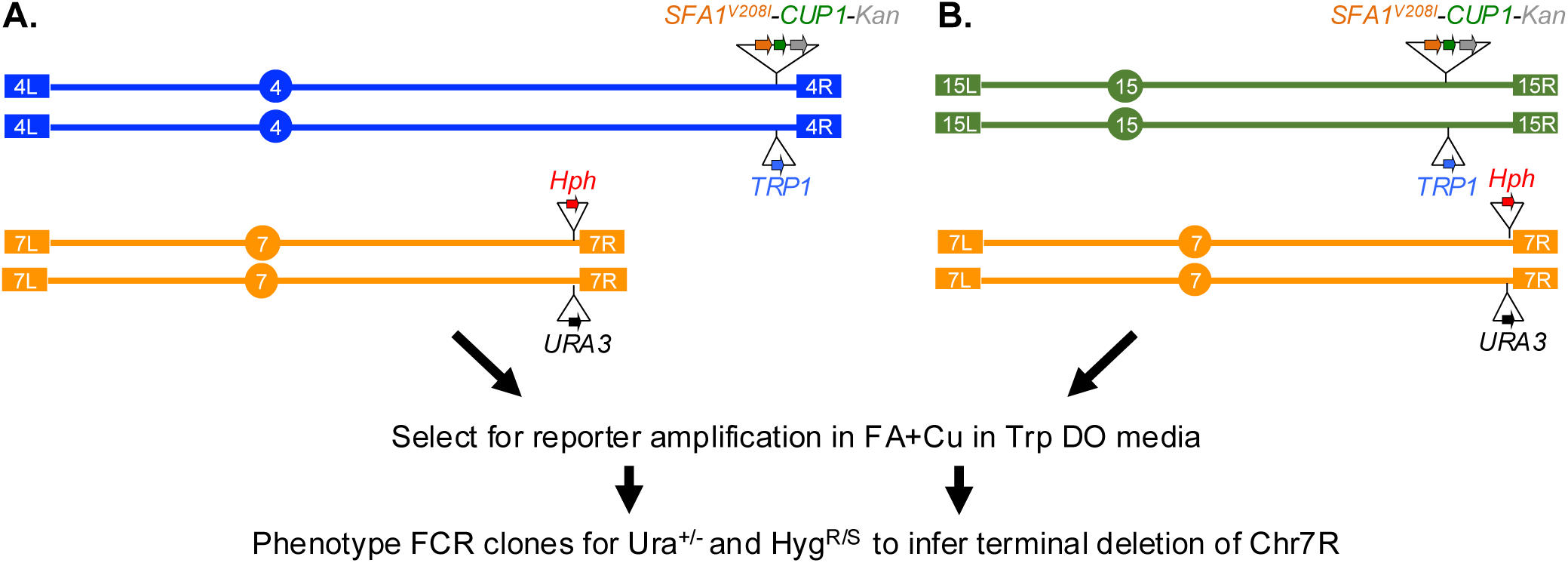
Phenotypic assessment of Chr7R deletion frequency in FCR clones selected for carrying amplifications on Chr4R or Chr15R. In both Chr4R and Chr15R parent strains, Chr7R possesses hemizygous telomere-proximal insertions of *_Kl_URA3* or HphMX markers downstream of *MAL13*. These strains enabled selection for amplification of the respective chromosomal region where the *SFA1^V208I^-CUP1* reporter was inserted, followed by screening and phenotypic inference of a concurrent deletion of Chr7R in either Chr7 homolog.

We grew independent cultures of the parent strains with the Chr4R and Chr15R reporters in rich media, and then plated them on tryptophan drop-out media containing the concentrations of combined formaldehyde plus copper sulphate (FA+Cu) that would only allow for cells possessing two or more copies of the amplification reporter cassette to grow and form FCR colonies. After these clones were recovered and purified, their growth phenotypes were retested on the same level of FA+Cu they were initially recovered from, and finally tested separately on uracil drop-out and on hygromycin B-containing media to interrogate the loss of either terminal region of Chr7R. This phenotypic screening of FCR clones from the two parents validated the initial biased behavior we had detected previously using genomic analyses (Table 1). We detected Chr7R deletions in 126 of 205 screened FCRs derived from amplification of the Chr4R reporter (61%) and in 77 of 158 FCRs derived from amplification of the Chr15R reporter (49%). In both parent strains, deletions of each Chr7R homolog were detected at approximately equal frequencies (66 Ura^−^ Hyg^R^ and 60 Ura^+^ Hyg^S^ among the Chr4R FCRs; and 41 Ura^−^ Hyg^R^ and 36 Ura^+^ Hyg^S^ among the Chr15 FCRs), consistently with both Chr7 homologs behaving similarly in these otherwise homozygous diploids.

### Is frequent Chr7R deletion due to a FCR selection bias?

Toward investigating possible causes of the pronounced bias of Chr7R deletion accompanying amplifications of our reporter cassette from other chromosomes, we first inquired whether the presence of a Chr7R deletion may somehow confer an advantage to growth in media containing FA+Cu. If such a selection bias is somehow granted by loss of Chr7R, we would expect to observe a more vigorous growth phenotype of FCR clones carrying Chr7R deletions compared to those with alternative chromosome arms deleted. We tested the relative viability and tolerance to FA+Cu of multiple FCR clones selected for amplification of Chr4R or Chr15R, noting where deletions occurred in each clone. Three matched sets of FCRs were identified from our collection, all carrying non-reciprocal translocations in which exactly the same terminal chromosome regions were amplified but associated with different terminal deletions so that their effect could be isolated and evaluated (Fig. 3; Top, Middle, Bottom sets). Independently isolated pairs of FCRs carrying the same rearrangements (*e.g.* FCR155 and FCR355) were also included in these tests to interrogate the degree of variability of the FA+Cu resistance phenotype between clones with equivalent karyotype rearrangements. While phenotypic heterogeneity was observed, these assays did not reveal a discernable pattern of resistance advantage for any particular regional chromosomal deletion, and FCR growth exhibited as much variation across clones possessing deletion of the same chromosome as it did between deletions of different chromosomal regions. These results suggested that Chr7R deletion either does not inherently confer a selective advantage for growth on FA+Cu relative to other deletions, or if it does, this effect is not sufficiently strong to explain the high abundance of Chr7R deletions recovered among FCRs clones.

**Figure 3.**
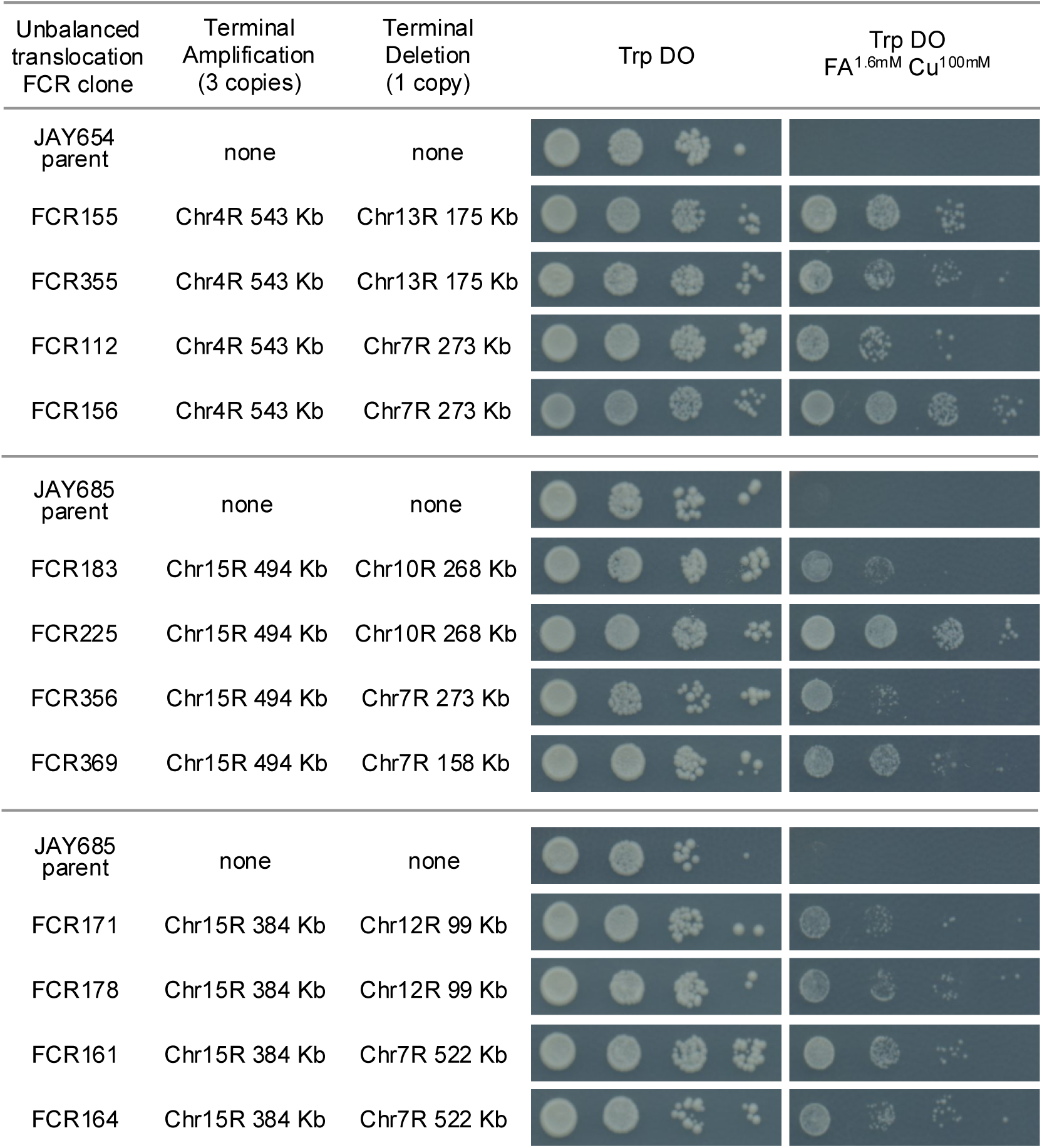
Comparison of formaldehyde and copper resistance phenotypes among FCR clones. The FCR clones shown all carried unbalanced (non-reciprocal) translocations, resulting in 3 copies of the amplified regions from Chr4R or Chr15R (2 copies with *SFA1^V208I^-CUP1*, 1 copy with *TRP1*), whereas their respective diploid parent strains carry the normal 2 copies of the same region (1 copy with *SFA1^V208I^-CUP1*, 1 copy with *TRP1*). Concurrently, each FCR clone had a terminal deletion (1 copy) of the indicated chromosome arms. The terminal amplification and terminal deletion tract lengths are indicated (*i.e.*, distances based on the CNV endpoints determined by array-CGH). The panels to the right show growth of 10-fold serial cell dilutions spotted on 1.6 mM FA + 100 µM Cu in a tryptophan drop-out solid media base and on the matched control permissive tryptophan drop-out without FA or Cu. The parent strains are sensitive to this concentration combination of the FA+Cu, while all FCRs display variable levels of resistance to FA+Cu, irrespective of the chromosomal deletion region.

### Investigation of possible Chr7R fragility

Next, we investigated whether this strong deletion bias might be caused by a high frequency of spontaneous DNA breakage somewhere along Chr7R, for example, through the presence of a discrete fragile site (Lemoine et al. 2005; Tang et al. 2011). For that purpose, we created a hybrid diploid strain by crossing a CG379-isogenic haploid to a haploid isogenic with the diverged YJM789 strain background (Wei et al. 2007), using an approach routinely used to map mitotic recombination tracts (St Charles and Petes 2013; Sampaio et al. 2020; Stewart et al. 2021). This diploid is heterozygous for thousands of single nucleotide polymorphisms evenly distributed genome-wide, including along Chr7R (Fig. 4A), which can be followed by microarray (array-SNP)(Zhang et al. 2013) or short read whole genome sequencing (WGS) genotyping (Heasley et al. 2021). We inserted the *_Kl_URA3*-*_Sc_URA3*-Kan CORE2 cassette at a position distal to the *MAL13* gene in the CG379 Chr7 homolog, approximately 18 Kb from the right telomere (*TEL07R*). This setup allowed us to select for clones that became resistant to 5-FOA after losing function of the counter selectable double *URA3* markers. All of such spontaneous clones concomitantly lost the Kan G418 resistance marker that is also present in the CORE2 cassette, thus confirming regional loss-of-heterozygosity (LOH), most likely due to allelic interhomolog mitotic recombination triggered by a break lesion in the CG379 Chr7R somewhere in 575 Kb between the centromere (*CEN7*) and the CORE2 insertion. This mechanism was confirmed through mapping the SNP markers that showed heterozygosity on the full left arm and up to discrete recombination sites on the right arm. A transition to homozygosity for the YJM789 SNPs was then used to infer the general vicinity of the precursor DSB lesion (Fig. 4B). If recombination in the assay region initiates primarily as a result of random DSBs, then the LOH tract endpoints should be distributed evenly. In contrast, the presence of a strong fragile site should yield a pattern of recurrent LOH endpoints clustered around a discrete region. We isolated 59 independent 5-FOA^R^ G418^S^ clones and mapped their LOH tracts using SNP-arrays or WGS (17 and 42 clones, respectively; Table S1). All clones had copy-neutral LOH tracts (loss of the CG379 SNPs and doubling of the YJM789 SNPs), and their endpoints were evenly scattered along Chr7R (Fig. 4B). We did not observe any LOH endpoint clustering pattern, including in the vicinity of a pair of double Ty element insertions present in Chr7R (corresponding to deletion endpoint sites *b* and *d* from Fig. 1). Similar structural tandem direct or inverted configurations of repetitive Ty1 and Ty2 sequences have previously been shown to have fragility properties and promote recombination on Chr3R under replication stress conditions (Lemoine et al. 2005), yet these did not appear to trigger excessive allelic mitotic recombination on Chr7R under the normal growth conditions we selected LOH clones from.

**Figure 4.**
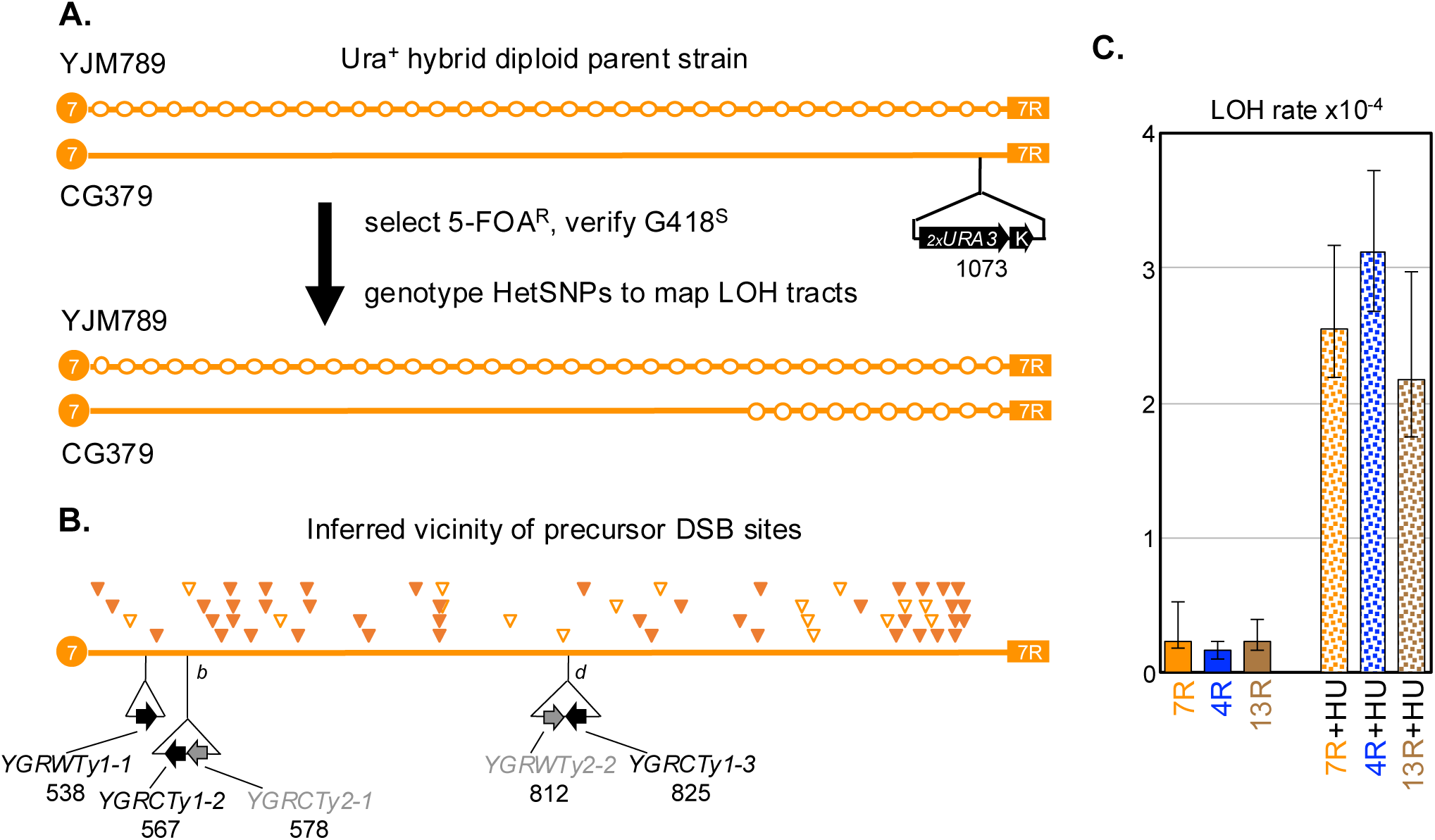
Investigation of Chr7R fragility. A. Hybrid diploid strain made by crossing a CG379-isogenic haploid to a haploid isogenic with the diverged YJM789 strain background. This strain possesses the *_Kl_URA3*-*_Sc_URA3*-Kan CORE2 cassette inserted distal to *MAL13* on Chr7R, which allowed for selection of LOH events on most of that arm. The filled circle at the left end represents the centromere (*CEN7*) and the rectangle at the right end represents the right telomere (*TEL07R*). Empty circles illustrate heterozygous SNP marker sites YJM789 differs from CG379 (not drawn to scale or number). B. LOH endpoints (inverted triangles) mark the beginning of the mapped YJM789 homozygosity tracts, and the inferred vicinities of precursor DSB sites in 59 Chr7R LOH clones (WGS [filled]; SNP-array [empty]; coordinates in Table S2). The positions (in Kb) of all full length Ty1 and Ty2 insertions (arrows) present on the CG379 Chr7R homolog are shown for reference relative to the mapped LOH endpoint sites. *b* and *d* correspond to the deletion endpoint sites *b* and *d* displayed in Fig. 1. C. Comparisons of LOH rates (LOH/cell/cell division) for Chr7R, Chr4R, and Chr13R with and without exposure to 75 mM HU.

This experimental system also gave us an opportunity to determine whether the rate of LOH in Chr7R might be higher than at comparably marked regions of the *S. cerevisiae* genome. We created similar CG379 x YJM789 hybrid diploids carrying a CORE2 cassette insertion on either the right arms of Chr4 or Chr13, in each case approximately 640 Kb distal from their respective centromeres, a distance ∼10% longer than in the Chr7R hybrid diploid described above. We used these three strains in quantitative fluctuation assays to directly measure and compare their LOH rates. All three strains had similar LOH rates when allowed to accumulate mutations spontaneously (Fig. 4C). We also measured LOH rates after growth in the presence of 75 mM HU prior to plating, a condition expected to induce fragile site activity through DNA replication stress (Arlt et al. 2012). While HU exposure resulted in 10- to 17-fold increase in the LOH rate in each chromosomal region, that increase was of comparable magnitude between them. Taken together, the random positional distribution of LOH endpoints along Chr7R, and the similar LOH rates relative to Chr4R and Chr13R both minus and plus HU exposure, disfavored the model where Chr7R contains a fragile site strong enough to explain the deletion bias detected among the FCR clones isolated earlier (Table 1; Fig. 2 and associated text).

### Orthologous validation of the Chr7R NAHR bias

We sought to develop a second, orthologous experimental approach that could independently recapitulate our initial observations made in FCR clones selected for amplification of the *SFA1^V208I^-CUP1* reporter that suggested a biased NAHR behavior of the Chr7R region. We adapted the conventional truncated-overlapping selectable marker approach widely used in *S. cerevisiae* recombination studies. In this case, we created a series of diploid strains homozygous for a 3’-truncated allele of the *URA3* gene (*URA*) at is native locus on the left arm of Chr5 (Chr5L). We then integrated a cassette containing 5’-truncated alleles of *URA3* (Kan-*RA3*) at seven test regions of the genome, including Chr7R (Fig. 5A). NAHR between the Chr5L *URA* and the Kan-*RA3*s is mediated by the 623 bp central *RA* shared homologous sequence and leads to the formation of a functional *URA3* full-length gene at the junction of a translocation between Chr5L and the respective Kan-*RA3* insertion chromosome. In all cases, the Kan-*RA3* cassette was integrated at sites similarly distant from their nearest telomere (246 to 257 Kb range), and all where relatively far from their respective centromeres (306 to 834 Kb range). The orientation of all *RA3* insertions was set to ensure the formation of viable monocentric translocations, and their positions, all in long chromosome arms, were chosen to support unconstrained mobility of the *RA3* substrates in the “Rabl-like” spatial arrangement of the yeast genome (Therizols et al. 2010; Agmon et al. 2013).

**Figure 5.**
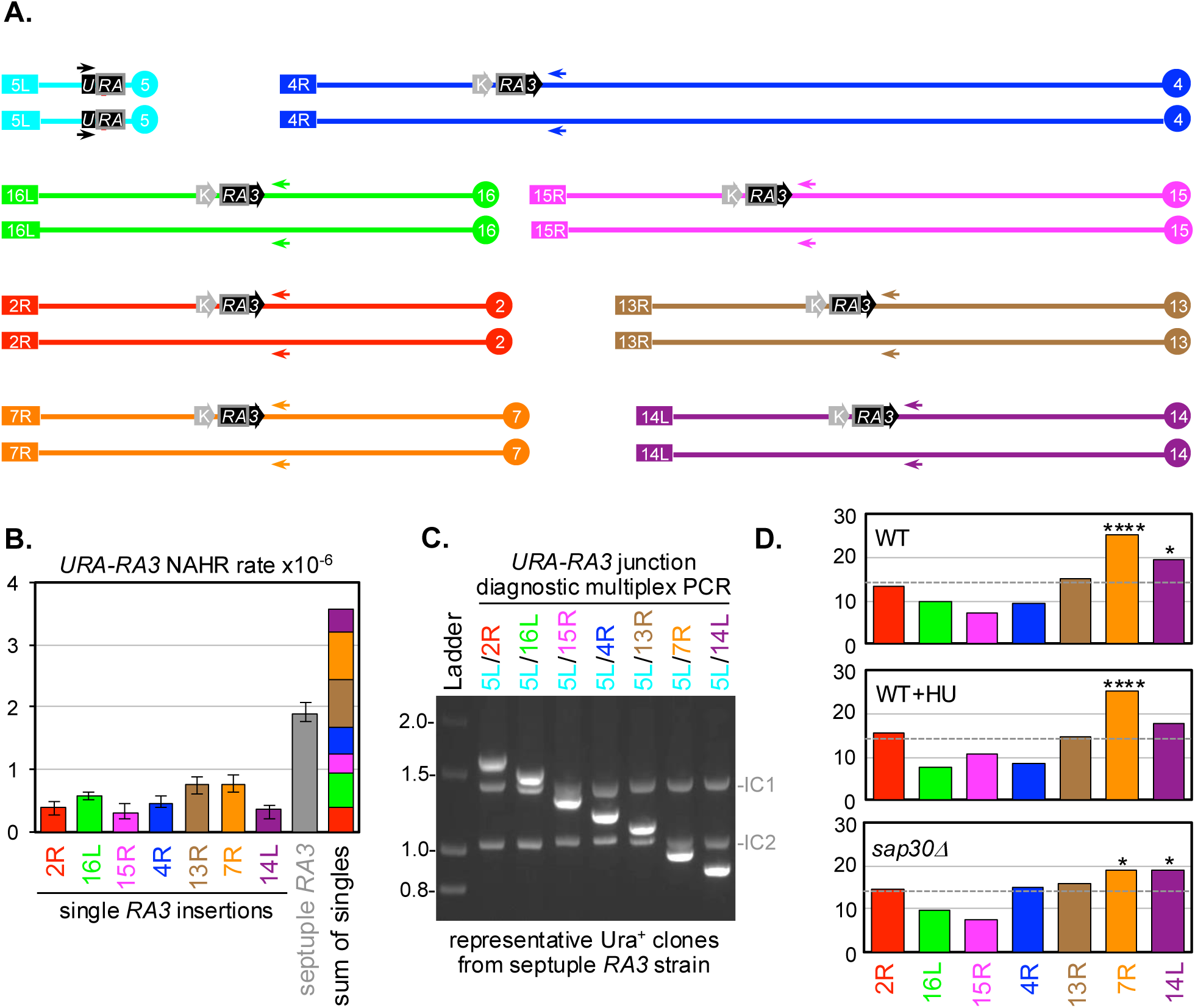
Orthologous validation of the Chr7R deletion bias: NAHR competition assay. A. Schematic representation of the site of the 3’-truncated allele of *URA3* gene at is native locus on the left arm of Chr5 (*URA*), and the integration sites of the cassette containing 5’-truncated alleles of *URA3* (Kan-*RA3*) at seven test regions of the genome. Only the relevant chromosome arm (centromere to telomere) for each test chromosome is shown for simplicity. Black box arrows represent the *URA* and *RA3* recombination substrates. The gray K block arrow represents Kan. Color-matched arrowheads represent the primers used in the multiplex PCR assay shown in C and D, with their respective orientations and positions flanking the *RA3* recombination substrates. B. *URA-RA3* NAHR rates leading to Ura+ colonies, for strains with each of the single *RA3* insertions and also for the septuple *RA3*-insertion strain. The multi-color column to the right represents the sum of all NAHR rates for the single *RA3* insertions. C. Agarose gel electrophoresis showing the multiplex PCR approach for detecting *URA-RA3* junctions from rearrangements involving the various *RA3* substrates on different chromosome arms. The fainter bands within the lanes correspond to the PCR products of two positive internal amplification sizing controls (IC1 and IC2), derived from genomic regions not involved in NAHR events. D. Results of the NAHR competition assay, showing the relative proportions of each *RA3* site involvement in NAHR events in the septuple *RA3* insertion strain. Y-axis indicates the percentage (%) across all Ura+ events analyzed by multiplex PCR; the dashed line indicates the neutral/random *RA3* utilization proportion (1 in 7; or 14.3%). Upper plot WT; wild type septuple *RA3*-insertion diploid; Middle plot WT+HU; wild type septuple *RA3* diploid exposed to HU 75 mM; Lower plot *sap30Δ/sap30Δ* septuple *RA3* diploid. **** *p*<0.0001 for *RA3* enrichment. * 0.05<*p*<0.01 for *RA3* enrichment.

We then used this experimental system to select Ura^+^ colonies, counting them to calculate the NAHR rate associated with each of the chromosomal regions being tested. We initially measured NAHR in diploid strains carrying one Kan-*RA3* insertion at a time and compared their rates (Fig. 5B; left side single-colored columns). We found that the seven *RA3* insertions individually enabled NAHR rates within a narrow ∼2.6-fold range, with the *RA3* at Chr7R among the ones most prone to recombination (albeit not-significantly higher). As a frame of reference, we also created a diploid with a Kan*-RA3* insertion at the right arm of Chr5, 253 Kb from *TEL05R* (not drawn in Fig. 5A). This strain, in which Chr5R *RA3* was physically tethered to one of the Chr5L *URA* substrates, had an NAHR rate ∼24-fold higher than the average *RA3* inserted at the other seven chromosome arms (12.1×10^−6^; 11.0-17.4×10^−6^ 95%CI; tethered rate not displayed in Fig. 5B). This was consistent with previous NAHR work in *S. cerevisiae* and validated the expectation that facilitated contact between *RA* substrates should lead to more frequent recombination in our system.

Next, we used the constructs above to create a new strain in which, instead of comparing the recombination properties of the seven *RA3* insertions individually, we could now have them all simultaneously available in the same genome and thus pitched in direct competition for engagement with the *URA*s on Chr5L. We conducted multiple rounds of crosses between haploids carrying individual *RA3s*, followed by meiosis, tetrad dissection, and selection, until we obtained recombinant haploids of opposite mating types containing triple and quadruple Kan-*RA3* insertions. These were then mated to form the diploid shown in Fig. 5A that contained all seven *RA3* insertions, one at each of the test chromosomal regions. We measured the NAHR rate in this septuple strain and found that it was 53% lower than the sum of the seven individual rates (Fig. 5B; right side; dark gray column compared to multicolor column). This suggested that in the septuple strain, the maximum NAHR potential was not fully realized, possibly due of the ability of specific *RA3* insertions to take priority over others and more avidly recombine with *URA*. If such a dynamic NAHR competition scenario does exist, then the expectation is that a qualitative analysis of the recombination products present among Ura^+^ clones should uncover a non-random distribution of *RA3* utilization.

In order to facilitate the characterization of large numbers of Ura^+^ recombinants, we developed a straightforward multiplex PCR approach (Fig. 5C). We designed a series of seven reverse primers, unique to each of the competing chromosomes, annealing to positions centromere-proximal of their respective Kan-*RA3* insertions at increasing nucleotide distances. A forward primer was designed to anneal at a fixed position of the *U* region of the Chr5L *URA* recombination substrates. PCR products running across the *URA3* translocation junctions had discernable lengths specific to the chromosomes that recombined with Chr5L. Two additional primer pairs were designed to generate internal control bands, amplified from centromere-proximal regions of two chromosomes not involved in the competition (Chr10 *CYR1* and Chr9 *PAN1*). All twelve primers were combined for multiplex PCR reactions using as template the genomic DNA from individual Ura^+^ clones. The amplification products were run on agarose gels, and the specific translocation product size detected in each Ura^+^ template was used as a diagnostic of which of the seven possible *RA3*s was present at the respective translocation junction.

We isolated 260 independent Ura^+^ clones derived from the septuple Kan-*RA3* diploid. Multiplex PCR of 249 of these provided unambiguous identification of the chromosomes involved in their NAHR events (Fig. 5D, upper plot; 11 Ura^+^ recombinants did not amplify any products; Table S2). The involvement of the tested chromosomes was significantly different (*p*<0.0001; Chi Square Goodness of Fit Test) from a neutral model prediction where each *RA3* contributes equally and randomly toward the total translocations (1 in 7; 14.3% null frequency). Remarkably, the distribution was strongly skewed toward a preference for the specific *RA3* inserted at Chr7R (*p*=4.5×10^−7^), and only mildly enriched for the *RA3* inserted at Chr14L (*p*=1.3×10^−2^). Next, we asked whether this biased distribution in the NAHR competition assay might be related to a frequent spontaneous breakage near the *RA3* inserted at Chr7R (*i.e.*, fragile site scenario). If that were the case, we reasoned that introduction of high levels of random genome-wide damage through replication stress might attenuate the differences in usage between *RA3*s and thus flatten-out the distribution. We grew the septuple *RA3* cells in the presence of 75 mM of HU, selected 200 independent Ura^+^ clones, and obtained 179 unambiguous PCR translocation junction calls (Fig. 5D, middle plot; 1 double *RA3* call, plus 20 without any amplification; Table S2). As expected, HU exposure led to a robust ∼5-fold increase in the rate of NAHR (data not shown), however, their qualitative distribution remained non-random (*p*=2.8×10^−5^), and importantly, not statistically different from the *RA3* usage distribution obtained from spontaneous conditions (*p*=0.5451; Kolmogorov-Smirnov Test), with the *RA3* inserted Chr7R continuing to be the clear competition winner (*p*=2.8×10^−5^).

Taken together, the results of the *URA-RA3* competition assay independently recapitulated the phenomenon of biased NAHR behavior associated with the Chr7R region, and supported the conclusion that this bias is not due to unusually high recombination initiation at that arm (*i.e.*, chromosome fragility). Instead, our observations suggest an alternative model that, upon breakage, Chr7R sequences display an inherently low fidelity of allelic HR partner choice (*i.e.*, sister chromatid repair) compared to other regions of the genome, thus are more likely to engage with a non-allelic homologous donor sequence present elsewhere.

### Flipping the role of Chr7R in NAHR from recipient to donor

To further characterize Chr7R’s behavior in inter-chromosomal translocations, we moved the *SFA1^V208I^-CUP1* reporter to Chr7R itself, now requiring cells to acquire amplifications of Chr7R, rather than deletion, in order to survive on FA+Cu media and be recovered as FCR clones (Fig. 6A). Assuming the absence of a recombination directionality bias, Chr7R should be just as capable of serving as a translocation donor as Chr4R or Chr15R, which are chromosome arms of comparable in size and repetitive DNA content. Further, Chr4R and Chr15R were often recovered as Chr7R’s partners in translocations such as those represented in Fig. 1. Therefore, we would expect to recover FCR clones at high rates, and the majority of which should carry inter-chromosomal non-reciprocal translocations resulting in amplification of Chr7R associated with concurrent deletions of other chromosome arms, including Chr4R and Chr15R.

**Figure 6:**
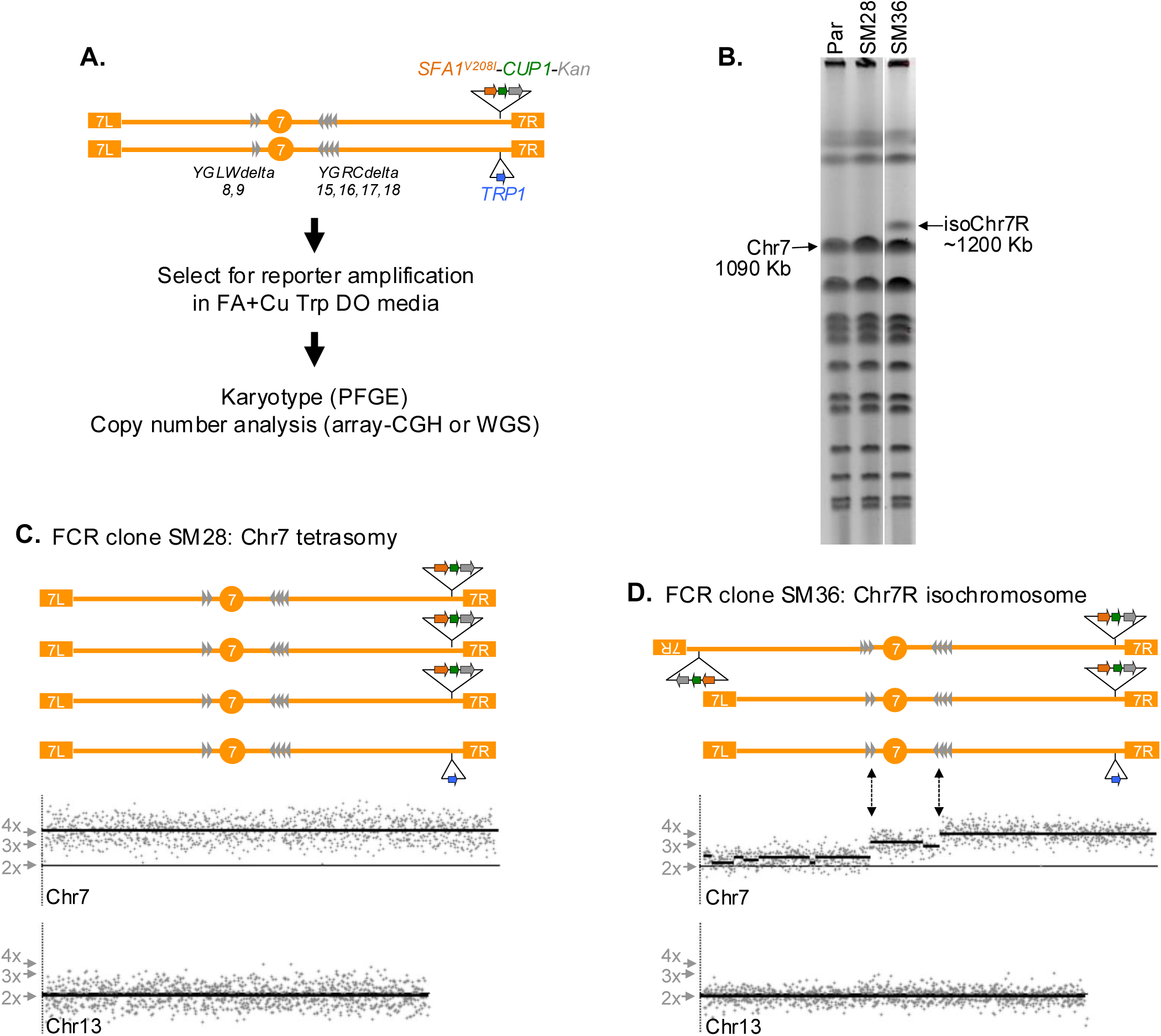
Karyotype analysis of FCR clones derived from amplification of the Chr7R region. A. Schematic representation of the experimental diploid strain with the *SFA1^V208I^-CUP1* reporter inserted on Chr7R, and workflow of the FCR selection and genomic analyses. Gray triangles indicate the approximate positions and orientations of delta LTR repeats present on either side of *CEN7*. These were the repeats involved in the rearrangement present in SM36, but note that many additional repetitive sequences, including full-length Ty elements, are present in Chr7 but are not shown in the figure for simplicity. B. Examples of the two categories of PFGE karyotypes observed in FCR clones derived from the parent Chr7R reporter strain shown in A (Par.; left lane). The SM28 FCR clone represents the category of whole chromosome gains which do not manifest as novel-sized rearrangement products, but a higher intensity band is seen at the Chr7 position. The SM36 FCR clone represents the category of intrachromosomal rearrangements, exhibiting a new ∼1200 Kb band associated with an isoChr7R. C. Array-CGH data showing the whole chromosome gain of Chr7R to four total copies. Upper section: schematic representation of the tetrasomic karyotype in SM28. Middle section: array-CGH copy number plot for Chr7, aligned under the Chr7 map with approximately matched coordinates. Lower section: array-CGH copy number plot for Chr13, showing a normal (2 copies) pattern for reference. No copy number alterations were detected at any other chromosomes in SM28. D. Array-CGH data showing segmented copy number increase across Chr7 associated with an intra-chromosomal rearrangement. Upper section: schematic representation of the isochromosome containing karyotype of SM36, plus the two normal Chr7 homologs still present. Middle section: array-CGH copy number plot for Chr7, aligned under the Chr7 map with approximately matched coordinates. Note the isochromosome configuration lacking the normal left arm segment (*TEL07L* to *YGLWdelta8-9*), but replaced with an inverted copy to the right arm segment that includes the amplification reporter (*YGLCdelta15-18* to *TEL07R*). Lower section: array-CGH copy number plot for Chr13, showing a normal (2 copies) pattern for reference. No copy number alterations were detected at any other chromosomes in SM36.

We integrated the reporter approximately 33 kb from *TEL07R*, between *YOR1* and *BGL2*. This site was chosen in part because it is close enough to the telomere to detect amplifications of a wide size range, with endpoints anywhere in the ∼560 Kb window between *CEN7* and *YOR1*. In addition, this site was chosen because it is distal to the last annotated Ty repeat (*YGRWdelta32*) and tRNA (*YNCG0046W*) sequences on Chr7R, but not within the subtelomeric or telomeric sequences. This location should favor translocation-type NAHR mediated rearrangements and avoid recovery of tandem segmental amplifications mediated by NAHR between Ty, tRNAs, or subtelomeric repeats. An insertion of the *TRP1* auxiliary selection marker was made at the corresponding allelic site on the other homolog of Chr7 present in the final diploid parent strain (Fig. 6A).

We carried out the FA+Cu resistance selection from this Chr7R parent strain and analyzed the genomes of 45 independently obtained FCR clones using a combination of PFGE, array-CGH, and short read WGS depth-of-coverage. Surprisingly, the copy number profiles derived from the analysis of these FCRs showed that amplifications of the *SFA1^V208I^-CUP1* reporter cassette were gained mostly through gain of extra copies of the whole Chr7 (trisomies and tetrasomies) or intra-chromosomal rearrangements, often involving the left arm of Chr7 (Table 2; Table S3). The analyses of FCR clones representative of these two predominant classes are shown in Fig. 6B-D. For example, FCR clone SM28 did not display any chromosome-size polymorphisms in PFGE (Fig. 6B), but had an array-CGH copy number profile consistent with 4 full-length copies of Chr7, 3 of which carried the *SFA1^V208I^-CUP1* amplification reporter (Fig. 6C). The intra-chromosomal rearrangement class is represented by FCR clone SM36, which displayed a new chromosomal band of ∼1,200 kb (Fig. 6B). Its copy number profile resembled a staircase climbing steps from left to right, with 2 copies from *TEL07L* to a region containing two Watson-oriented LTR repeats (*YGLWdelta8* and *YGLWdelta8*), followed by 3 copies of the segment extending through *CEN7* until the site of a double Ty insertion containing four Crick-oriented LTRs (*YGRCdelta15* to *YGRCdelta19*), and finally four copies of the right terminal region including the amplification *SFA1^V208I^-CUP1* reporter up to *TEL07R*. This CNV pattern and the ∼1200 kb PFGE band, are consistent with the presence of an isochromosome (isoChr7R) formed by NAHR between the LTRs present at the two copy-number transition endpoints, resulting in deletion of left arm and duplication of the right arm in reverse orientation (mirror image-like chromosomal molecule). These data also showed that, in addition to the isoChr7R intra-chromosomal rearrangement, SM36 retained its two original copies of Chr7 for a total three copies of the amplification reporter cassette.

**Table 2.** Distribution of classes of structural rearrangements among FCR clones selected for amplification of the *SFA1^V208I^-CUP1* reporter inserted at various chromosomal sites.

| Classes of structural genomic rearrangements leading to FA+Cu resistance | Number of FCR clones derived from strains with the respective *SFA1<sup>V208I</sup>-CUP1* reporter insertion sites and genotypes |  |  |  |  |  |
| --- | --- | --- | --- | --- | --- | --- |
|  | <sup>a</sup> Chr4R | <sup>a</sup> Chr15R | <sup>b</sup> Chr7R |  |  |  |
|  | WT | WT | WT | *swr1Δ* | *sae2Δ* | *sap30Δ* |
| Whole Chr reporter amplification<br>*i.e.* trisomy or tetrasomy | 5 | 1 | 28 | 13 | 3 | 28 |
| Intra-chromosomal reporter amplification<br>*i.e.* Chr7L deletion; isochromosome | 0 | 1 | 9 | 1 | 2 | 4 |
| Inter-chromosomal reporter amplification<br>*i.e.* non-reciprocal translocation | 20 | 22 | 4 | 3 | 0 | 10 |
| <sup>c</sup> Other (non-amplification) | 0 | 0 | 7 | 2 | 13 | 7 |
| Total FCR clones analyzed | 25 | 24 | 45 | 18 | 18 | 49 |

<sup>a</sup> FCR clones derived from Chr4R and Chr15R parent strain grown under uninduced conditions. Assessed based on array-CGH data from Stanton 2012.
<sup>b</sup> Individual FCR clone CNV data assessment (array-CGH and WGS) available in Table S3.
<sup>c</sup> FCR clones that did retest positive for resistance to FA+Cu, yet did not display a CNV signal anywhere in their genomes, including no sign of amplification in or near the *SFA1<sup>V208I</sup>-CUP1* reporter insertion sites. The FA+Cu resistance mechanism was not characterized in these clones.

The pattern of amplifications obtained when the *SFA1^V208I^-CUP1* reporter was inserted at Chr7R, dominated by whole chromosome gains and intra-chromosomal rearrangements, was in marked contrast to the pattern of FCR classes derived when the reporter insertion was on Chr4R or Chr15R, which instead were characterized by an abundance of inter-chromosomal non-reciprocal translocations (most of which involved a deletion on Chr7R). Only a relatively small subset of the Chr7R-derived FCRs (4/45) had terminal amplifications acquired through inter-chromosomal non-reciprocal translocations mediated by NAHR (Table 2). These results showed a substantial shift in the amplification patterns, likely associated with the unusual recombination behavior of the Chr7R region, which acted frequently as an inter-chromosomal NAHR translocation recipient, but rarely as a donor (inter-chromosomal translocations of Chr7R reporter vs. Chr4R reporter: *p*=2.3×10^−9^; inter-chromosomal translocations of Chr7R reporter vs. Chr15R reporter: *p*= 5.9×10^−12^; Fisher’s exact test).

Importantly, the Chr7R reporter strain (Fig. 6A) produced a substantially and noticeably lower number of FCR clones compared to the Chr4R and Chr15R experimental strains, so much so that we were unable to directly measure the rate of Chr7R reporter amplification. In addition, ∼15% (7 of 45) of these rarer Chr7R-derived FCRs did not contain any amplification of the genomic region containing the *SFA1^V208I^-CUP1* reporter, or any copy number alterations at other genomic regions either. Yet, these non-amplification FCRs clones did display a clear and stable FA+Cu resistance phenotype upon post-purification retest. This unexpected FCR class (Other / non-amplification in Table 2) was not detected among FCRs derived from the Chr4R and Chr15R experimental strains (Stanton 2012; Sharif 2014). While we have not yet characterized their alternative mechanism(s) of FA+Cu resistance, we interpret their detection in the Chr7R reporter strain as an indirect consequence of a low rate of reporter amplifications mediated by NAHR. Relatedly, this is also consistent with the relatively high incidence (∼62%; 28 of 45) of the NAHR-independent whole chromosome gain FCR class, which was rare among Chr4R- and Chr15R-derived FCRs. Assuming the rate of whole chromosome gain is generally similar between these three large *S. cerevisiae* chromosomes (Chr4, ∼1.5 Mb; Chr7 and Chr15 ∼1.1 Mb), the proportion of recovered FCRs caused by their tri- or tetrasomies might be an indirect reflection of their respective and variable NAHR rates. It follows that the unique distribution of FCR classes that characterizes the Chr7R reporter strain must be a consequence of this region’s reluctance to play the donor role in NAHR, in stark contrast to its eagerness to play the recipient role when the *SFA1^V208I^-CUP1* is inserted elsewhere (Table 1).

### Investigation of the potential involvement of DSB mobility and sister chromatid cohesion

We hypothesized two different models to explain Chr7R’s biased behavior in NAHR translocations, both related to a differential in spatial accessibility of non-allelic HR substrates between Chr7R and other regions of the genome. In one scenario, this behavior could be related to Chr7R possessing enhanced DSB mobility relative to other chromosomes, and another related to inherently weaker sister chromatid cohesion at Chr7R. Enhanced DSB mobility could allow Chr7R DSBs to explore a higher volume of the nucleus and thus more often engage in far-reaching homology search. Once broken, this type of mechanism might provide the Chr7R region with a greater efficiency in engaging distant ectopic donor HR substrates, thus accounting for its unusual propensity for serving as a translocation recipient (Table 1). Alternatively, sister chromatid cohesion at Chr7R could somehow be less effective than at other chromosomes, possibly due to faulty or diminished cohesin loading along part or all its length. In this case, DSBs within Chr7R would be less likely to be repaired accurately using the allelic sister chromatid template and would be more prone to engaging a distant ectopic template instead. Either one, or a combination of these models, could account for Chr7R’s tendency to frequently function as a translocation recipient. To begin exploring these models, we compiled a list of nine candidate genes whose deletions are viable and were previously shown to be involved in either DSB mobility or sister chromatid cohesion.

For investigating DSB mobility, we knocked out *RAD54*, *INO80*, *HTZ1*, *SWR1*, *SAE2*, and *RAD9*. Rad54 increases mobility of damaged loci through its ATPase activity and may be more important for inter-chromosomal homology searches than intra-chromosomal searches. Its role has been tested in haploid yeast, but not diploids (Mine-Hattab and Rothstein 2012). Ino80 is a subunit of the INO80 chromatin remodeling complex, which enhances mobility of chromatin sites it is bound to. This activity is dependent on its Ino80 ATPase subunit (Neumann et al. 2012). H2A.Z is a histone variant incorporated at DSB sites. Deletion of its encoding gene, *HTZ1*, was found to decrease mobility of a DSB site, but still allow localization of the site to the nuclear periphery (Horigome et al. 2014). Swr1 is a component of the chromatin remodeling complex SWR1, which replaces the H2A/H2B dimer of nucleosomes with the variant histone H2A.Z/H2B dimer. Like *HTZ1*, it was reported that deletion of the *SWR1* gene reduces DSB mobility while preserving DSB relocalization (Horigome et al. 2014). Sae2 is a nonessential protein which acts upstream of Rad51 in resection of DSBs, and it was previously reported that pairing and mobility of DSB loci were delayed but not prevented by deletion of *SAE2* (Mine-Hattab and Rothstein 2012). Finally, Rad9 is a non-essential DNA damage checkpoint protein. Deletion of *RAD9* has been shown to delay appearance of recombination intermediates which form during repair of DSBs and is implicated in long-range homology searches (Dion et al. 2012).

To probe a role for sister chromatid cohesion in the establishment of the Chr7R NAHR biased behavior, we selected three deletion-viable candidate genes: *TOF1*, *SAP30*, and *HDA1*. The Tof1 protein is part of a complex which promotes sister chromatid cohesion at stalled replication forks and facilitates their repair (Mayer et al. 2004). Similar to S phase degradation of the acetyltransferase Eco1, it was previously found that *tof1Δ* enhanced mobility of spontaneous S phase damage sites (Dion et al. 2013). Sap30 is a component of the Rpd3L histone deacetylase complex. *SAP30* deletion significantly decreases sister chromatid recombination and in turn increases ectopic recombination in repair of both induced and spontaneous DSBs which can arise during replication (Ortega et al. 2019). Hda1 is a subunit of the HDA1 histone deacetylase complex. As with deletion of *SAP30*, Ortega et al. found that its deletion significantly decreased sister chromatid recombination in repair of induced or spontaneous DSBs during replication; however, *HDA1* deletion did not increase ectopic recombination (Ortega et al. 2019).

We conducted initial screening tests to ask whether knocking out any of these candidate genes could eliminate or at least attenuate Chr7R’s strongly biased recombination behavior. To do this we built homozygous candidate gene deletions in the same diploid background used earlier in our study (Fig. 2A) to phenotypically quantify the frequency of Chr7R deletion (Ura^−^ or Hyg^S^) among FCRs derived from amplification reporter inserted on Chr4R. Three of the initial candidate deletions either had extremely slow growth (*INO80*, *HTZ1*) or did not support sufficient numbers of FCR clones to be recovered (*RAD54*), thus were not pursued further. FCRs derived from the six remaining candidate gene deletions were successfully isolated and scored for the frequency of Chr7R deletions relative to wild type (Table 3). Notably, in this round of this assay the frequency of Chr7R deletion among WT FCRs was even higher (89%) than we had measured earlier (61%). The reason for this variation is not known, but frequencies of Chr7R deletions among the mutant-derived FCR clones were compared to the frequency measured in contemporaneously isolated WT-derived FCRs (89%). We selected and analyzed FCR clones for the WT and gene deletion strains concurrently and progressively, initially ∼15-30 clones from each genotype, and we continued to isolate and score additional FCRs from those that appeared to lower the Chr7R deletion frequency relative to WT. In the end, deletions of three of the candidate genes (*SWR1*, *SAE2*, and *SAP30*) significantly attenuated the Chr7R deletion bias seen in WT, and were thus selected for further analyses (*swr1Δ* vs. WT: *p*=4.2×10^− 4^; *sae2Δ* vs. WT: *p*=9.4×10^−3^; *sap30Δ* vs WT: *p*=1.7×10^−3^; Fisher’s exact test).

**Table 3.** Frequency of Chr7R deletion among FCR clones selected for amplification of the *SFA1^V208I^-CUP1* reporter inserted at Chr4R.

| Genotype | <sup>b</sup> WT | *swr1Δ* | *sae2Δ* | *rad9Δ* | *tof1Δ* | *sap30Δ* | *hda1Δ* |
| --- | --- | --- | --- | --- | --- | --- | --- |
| <sup>a</sup> FCR clones with a deletion on Chr7R (Ura<sup>-</sup> or Hyg<sup>S</sup>) | 55<br>(89%) | 26*<br>(58%) | 34*<br>(68%) | 18<br>(78%) | 33<br>(100%) | 21*<br>(60%) | 13<br>(93%) |
| Total FCR clones tested | 62 | 45 | 50 | 23 | 33 | 35 | 14 |

<sup>a</sup> Concomitant Chr7R deletion inferred through Ura<sup>-</sup> or Hyg<sup>S</sup> phenotypes as shown in Fig. 2A.
<sup>b</sup> Matched control WT FCR clone set, isolated in parallel and contemporaneously to the mutant-derived FCRs shown in Table 3.
\* indicates significant ( $p < 0.01$ ) attenuation Chr7R deletion proportion relative to the matched WT control set<sup>b</sup> above (Fisher's Exact Test).

We next asked whether any of these three genes affected Chr7R’s capacity to act as a translocation donor using the experimental strain possessing the *SFA1^V208I^-CUP1* reporter inserted on Chr7R (Fig. 6A). We initially isolated 18 FCRs derived from each of the *swr1Δ*, *sae2Δ* or *sap30Δ* mutant strains and carried out copy number analysis using short-read WGS depth-of-coverage to allow their qualitative categorization and comparison of their amplification classes relative to WT (Table 2). *SWR1* deletion did not appreciably change the pattern of amplifications, specifically because it retained a relatively high proportion of whole Chr7 amplifications (trisomy and tetrasomy class). Loss of *SAE2* led to a high proportion of FCRs that did not possess any detectable amplifications (Other class), thus were difficult to interpret relative to WT and the other genotypes. As mentioned above, it is not yet known how the 13 of 18 *sae2Δ* FCR clones acquired FA+Cu resistance, but we hypothesize that this may be related to the general recombination defect in this strain, thus enriching for otherwise rare mechanisms of FA+Cu resistance that are independent of *SFA1^V208I^-CUP1* amplification.

Finally, deletion of *SAP30* did appear to alter the WT amplification pattern by increasing the proportion of inter-chromosomal rearrangements and enhancing Chr7R’s role as a translocation donor. We detected this trend in the initial 18 *sap30Δ* FCRs analyzed, which prompted us to isolate and analyze by WGS an additional 31 *sap30Δ* FCRs to reach a more robust 49 total clone set (Table 2). We found that while the proportion of whole chromosome gains remained stable in *sap30Δ* FCRs relative to WT, the number of intra-chromosomal amplifications was reduced (4/49 vs. 9/45 in WT; *p*=0.1361; Fisher’s exact test), while the number of inter-chromosomal amplifications increased (10/49 vs. 4/45 in WT; *p*=0.1516; Fisher’s exact test). This mild shift from intra- to inter-chromosomal observed in the *sap30Δ* FCRs was in a direction that resembled the pattern that characterizes the WT FCR derived from the reporter insertions on Chr4R and Chr15R. Collectively, the *SAP30* deletion both lowered the frequency of Chr7R deletion accompanying amplification from Chr4R decreasing Chr7R ability to act as a recipient (Table 3), and also increased inter-chromosomal rearrangements improving the ability Chr7R acts as a translocation donor (Table 2).

Given the observations trends above, that deletion of *SAP30* was able to mildly weaken the biased behavior of Chr7R in NAHR clones derived from FA+Cu resistance selection, we sought to recapitulate and validate this result using the NAHR competition assay approach (Fig. 5). We deleted both copies of *SAP30* in the septuple insertion *RA3* diploid strain, and like we had done before, isolated a large number of Ura^+^ clones carrying *URA-RA3* recombination products to characterize the relative frequency of participation of each *RA3* insertion in NAHR. We isolated 221 independent Ura^+^ clones derived from the *sap30Δ* septuple insertion *RA3* diploid. Multiplex PCR analyses of 216 of these provided unambiguous identification of the chromosomes involved in their respective *URA-RA3* NAHR events (Fig. 5D, bottom plot; 3 double *RA3* calls, plus 2 without any amplification; Table S2). In this case, the overall distribution of *RA3* NAHR participation was still significantly different from a random expectation (*p*=8.0×10^−3^, null frequency 14%, Chi Square Goodness of Fit Test), but not as pronounced as the deviation measured in wild type (*p*=1.0×10^−7^). The frequency of participation of the *RA3* inserted at Chr7R was still relatively high, but other *RA3* insertions rose in prominence relative to wild type, leading to a less biased distribution. Chr7R’s *RA3* participation frequency in the NAHR competition in *sap30Δ* was only marginally significantly different from the random expectation at the 0.05 significance threshold (*p*=4.4×10^−2^), whereas in wild type that bias was far more evident (*p*=4.5×10^−7^).

In summary, the trends of results from our analyses of *sap30Δ* mutant diploids were consistent across the two FCR selection assays (Tables 2 and 3) and the *URA-RA3* NAHR competition assay (Fig. 5) in detecting an attenuation of the biased behavior of the Chr7R region relative to that observed in control wild type cells. Taken together, these data suggest that Sap30, while not sufficient, is likely a contributing factor, important to the full manifestation of the biased Chr7R behavior in NAHR, possibly through the role of the Rpd3L complex in the establishment or preservation of sister chromatic cohesion following DSB formation.

## DISCUSSION

We set out to investigate the bias of Chr7R taking part in NAHR as a frequent translocation recipient. In the process, we have shown that it can be reproduced through two independent recombination assays using different selection approaches, and that Chr7R’s peculiar behavior is most likely the result of a repair bias rather than an initiation bias involving recurrent fragile site breakage. We also discovered that Chr7R’s behavior is unusual both as a translocation donor as well as a recipient. Specifically, this region of the *S. cerevisiae* genome engages in inter-chromosomal translocations as a recipient more often than other chromosome arms, yet, it functions as a translocation donor less frequently relative to others. Altogether, these data suggest that Chr7R possesses unusual recombination behavior relative to other chromosome arms comparable in size and repetitive DNA content. While the mechanism underlying this biased behavior remains unknown, in this work we conducted an initial exploration of two candidate pathways that may be involved: spatial mobility of broken DNA and sister chromatid cohesion (SCC).

Chr7R’s peculiar behavior might be attributable to a difference in mobility or localization relative to other chromosomes (Haber 2018; Chiolo et al. 2025). Increased mobility would be expected to enhance a chromosome’s ability to participate in NAHR as a translocation recipient, as with Chr7R, and conversely may also decrease the chance of a broken DNA end engaging its allelic template in the sister chromatid to result in accurate homologous repair. We propose that different chromosomes may have differing mobility capacity in the 3D space of the nuclear matrix and that this heterogeneity could modulate the ability for dispersed repeats within them to engage one another and become recombination partners. In order for NAHR to occur, a damaged chromatid must fail to find and engage its identical allelic sister or the corresponding allelic site in the homologous chromosome, and instead it must find and engage a non-allelic repair template. Region-specific variation in the effectiveness of one or both of these key steps could potentially lead to differential repair outcomes at affected genomic segments. A regional defect or delay in SCC, might make that segment more vulnerable to engaging a non-allelic partner in an inappropriate repair event, leading to structural rearrangements.

The Sap30 protein is a subunit of the histone deacetylase complex (HDAC) Rpd3L, which has been shown to facilitate cohesin loading and associated SCC, promoting efficient sister chromatid recombination (Ortega et al. 2019). Without Sap30, the function of the complex is compromised, and the resultant reduction in cohesin loading leads to deficient SCC, indirectly leading to greater NAHR. It is not currently known which histones or histone residues Rpd3L deacetylates to promote loading of cohesin, nor by what mechanism the deacetylation of its target histones causes this effect (Gomez-Gonzalez et al. 2020). The relationship between HDAC activity and DNA repair has yet to be fully understood, as well as how HDAC activity affects SCC in human cells. Our study of these pathways in yeast offers an additional facet to the current understanding of the mechanisms by which HDAC inhibition leads to genomic instability, and possibly to its associated biomedical implications.

Deletion of either *SWR1*, *SAE2*, or *SAP30* lowered the percentage of FCR clones sustaining a concurrent Chr7R deletion accompanying amplification of Chr4R, although a pronounced NAHR bias remained. This suggests that all three of these proteins might differentially contribute to Chr7R’s ability to find homologous templates as a translocation recipient relative to other chromosomes, and their loss may somehow impair Chr7R’s mobility, or enhance the mobility of other chromosomes to attenuate the prevalence of Chr7R as a translocation recipient. This mechanism could explain why *SAP30* deletion decreases the number of Chr7R deletions accompanying *SFA1-CUP1* Chr4R amplifications in inter-chromosomal rearrangements and also attenuated the bias of Chr7R participation in the *URA-RA3* NAHR competition assay. In this scenario, *sap30Δ* mutant cells would have faulty SCC throughout the genome, which in turn could grant other chromosomes more opportunity to recombine with the Chr4R reporter arm. Likewise, *RA3* repeats present at regions other than Chr7R became more frequent partners in non-allelic recombination with the Chr5L *URA* substrate.

When FCR clones possessing amplifications of Chr7R were recovered, the proportion of inter-chromosomal rearrangements facilitating these amplifications was markedly lower than in clones possessing amplifications of Chr4R or Chr15R. We propose that another contributing factor to this bias could be Chr7R somehow having relatively a higher mobility which renders it an elusive target for damaged chromosomes to find or use as a repair template to receive translocations from. However, we observed a trend of increased inter-chromosomal rearrangements accompanying amplification of Chr7R when *SAP30* was deleted. Deletion of *SWR1* or *SAE2* did not cause such an increase. The increase in ectopic recombination associated with *SAP30* loss has been observed by others (Ortega et al. 2019). We speculate that *SAP30* deletion may enhance Chr7R’s function as a translocation donor by removing the natural mobility constraint of SCC for the damaged chromosomes which receive translocations from Chr7R rather than by affecting Chr7R itself. It is more likely that *SAP30* deletion affects the translocation recipient chromosomes in this case, because DSB sites experience a greater expansion in their mobility than do the intact template loci they use during repair processing (Dion et al. 2012; Mine-Hattab and Rothstein 2012). Therefore, there may be a greater range of movement to be affected in recipient chromosomes. In this scenario, *SAP30* deletion could allow these translocation recipients to more closely mimic Chr7R’s ectopic tendencies as a translocation recipient.

In future follow-up work, it could prove valuable to directly characterize the mobility properties of Chr7R comparatively to other regions of the genome, both before and after DSB induction (Hajjoul et al. 2013). The chromosome conformation capture approaches could be used to evaluate the relationship between chromosome mobility and the observed mutations in FCRs in the context of our diploid experimental strain system. This could allow comparisons of chromosomal contact frequencies which do and do not involve Chr7R, perhaps offering insight into whether Chr7R possesses a heightened mobility that facilitates NAHR. Fluorescent protein labeling and live-cell microscopy could also be used to visualize Chr7R’s movement in real time relative to another site in the genome known to be comparatively static.

Our study suggests that the contributions of HDACs and SCC is an aspect of structural variation genesis that warrants to further exploration. We have shown that loss of a single HDAC subunit can qualitatively modulate ectopic recombination preferences, but that the change does not affect all chromosomes equally. Further investigation of this bias in the yeast model could pave the way for better unraveling the mechanisms behind recurrent mitotic chromosomal rearrangement events more broadly in higher eukaryotes.

## MATERIALS AND METHODS

### Yeast growth conditions

Yeast strains were grown on rich YPD media containing 10 g yeast extract, 20 g glucose, 20 g peptone, and 20 g bacteriological agar in 1 L of distilled water. Geneticin (Gen), Hygromycin B (Hyg), or Nourseothricin (Nat) were supplemented to solid YPD as needed (Goldstein and McCusker 1999). Hydroxyurea (HU) was supplement at 75 mM to solid YPD. Synthetic drop-out (DO) media contained 1.7 g yeast nitrogen base without amino acids, 1.4 g of the appropriate DO mix (USBiological Life Sciences), 5 g ammonium sulfate, 20 g glucose, and 20 g bacteriological agar in 1 L of distilled water. 5-FOA (USBiological Life Sciences) was supplemented at 1 g/L concentration in synthetic media with complete dropout mix.

For FCR selection and resistance phenotyping, synthetic Trp DO media was supplemented with formaldehyde (FA) and CuSO4 (Cu). Given the labile nature of FA in diluted aqueous solution, FA was added to media from a fresh dilution of a 37% stock solution stabilized in 10% methanol (Sigma F-8775). A 1 M FA dilution from this stock was prepared fresh in sterile water and added to Trp DO agar media after autoclaving, and shortly before pouring Petri plates. Trp DO with FA+Cu was typically used within 24-48 hours for the selection of FCR clones. The specific FA+Cu concentration combination that inhibited growth of cells carrying one copy of the *SFA1^V208I^-CUP1* reporter, but allowed growth of cells carrying two or more copies of the reporter varied between experiments depending on the site of reporter insertion, strain genotype, and batch of 37% FA stock. Thus, the specific FA+Cu combination used in FCR selection was re-calibrated periodically using control strains with known copy numbers of the reporter as calibration standards. Cultures in solid media containing FA+Cu were incubated for up to 5 days at 30C, inside a closed plastic tub containing a damp paper towel to maintain humidity and prevent desiccation. Liquid YPD cultures were grown at 30 C overnight in a rotating drum for culture tubes.

### Yeast strain backgrounds and strain construction

All strains used in the selection of FCR clones were isogenic strain MS71, a close derivative of the CG379 strain background (Morrison et al. 1991; Argueso et al. 2008; Zhang et al. 2013). The parent strains with the *SFA1^V208I^-CUP1* reporter integrated at Chr4R, Chr5R, and Chr15R were previously described (Stanton 2012). The specific strains and all oligonucleotides used in this study are listed in Table S4 and Table S5, respectively.

All site-specific genetic modifications used in experimental diploid strains were built first into MS71-isogenic haploids, which were then mated to generate diploids hemizygous or homozygous at the respective sites. The *SFA1^V208I^-CUP1* reporter or the *TRP1* auxiliary marker were integrated between *YOR1* and *BGL2* near the right end of Chr7R. Strains carrying candidate gene deletions were created by the PCR-mediated approach with drug resistance markers, typically NatMX (Goldstein and McCusker 1999). The *Kluyveromyces lactis KlURA3* and HphMX markers were integrated between *MAL13* and *MAL11* near the right end of Chr7R. All marker integrations and gene deletions were verified by PCR using primers to validate both the upstream and downstream integration sites. The hybrid diploid strains used in the LOH rate and endpoint mapping experiments used as one of the haploid parents a strain isogenic with the YJM789 background described previously (Wei et al. 2007; Sampaio et al. 2020). The strains used in the *URA-RA3* NAHR competition assays were built by individual insertions of the KanMX-*RA3* cassette at seven genomic sites in MS71-isogenic haploids, followed by sequential rounds of crossings, tetrad dissection and meiotic recombinant spore selection. The specific genomic coordinates of integrations and deletions are provided in Table S5.

### Chr7R FCR selection

Chr7R FCR clones were identified as colonies growing on solid Trp DO media containing a concentration of FA+Cu empirically determined to be minimally sufficient to prevent the growth of their respective parent strain with one copy of the *SFA1^V208I^-CUP1* reporter. Relative to Chr4R, Chr5R, and Chr15R (Stanton 2012), the Chr7R *SFA1^V208I^-CUP1* reporter insertion described in this study resulted in lower frequencies of FCR colonies, and comparatively higher emergence of small background colonies displaying marginal FA+Cu resistance. These background colonies prevented the reliable quantitative measurement of FCR mutation rates, similarly to limitation described earlier for our original *SFA1-CUP1* reporter (Zhang et al. 2013). For each FCR clone isolated, a 5 mL YPD culture of its parent was first inoculated and allowed to grow for 24-48 hrs. 1 mL of each culture was spun down and then washed with two 1 mL rinses of sterile water. The washed cells were resuspended in another 1 mL of water and 150 µL of this suspension was plated on the appropriate FA+Cu concentration, and incubated at 30C for 5 days. A single FCR colony was isolated from each culture to ensure mutational independence, then streaked to single colonies in Trp DO with a lower concentration of FA+Cu, and then retested to validate their FCR phenotype through serial dilution spot assays on Trp DO with the higher FA+Cu concentration the respective clone was originally isolated from. FCRs derived from strains carrying the *MAL13::klURA3* / *MAL13::HphMX* hemizygous markers were subsequently tested on Ura DO and YPD plus Hyg to assess the presence of a concomitant deletion on Chr7R.

### *URA-RA3* NAHR and LOH clone selection and rate measurements

Single colonies from Ura^−^ parent strains carrying single or septuple insertions of the *RA3*-KanMX cassette were grown in liquid YDP overnight and appropriate dilutions were plated to selective Ura DO or permissive YPD. Colony counts from these plates were used to calculate the rate of *URA-RA3* NAHR using the method of the median and as described earlier (Lea and Coulson 1949; Hall et al. 2009; Conover et al. 2015). LOH rate measurements and clone selection was carried out as described earlier (Rodrigues-Prause et al. 2018). NAHR and LOH clones were isolated as a single colony per culture to ensure mutational independence, streaked to single colonies non-selectively, and the purified clones were validated phenotypically in Ura DO or 5-FOA before multiplex-PCR or SNP-array genomic characterization, respectively.

### Genomic analyses

Karyotyping by pulse-field gel electrophoresis (PFGE), Chr7R LOH endpoint mapping with SNP-array, copy number profiling by array-based comparative genomic hybridization (array-CGH and short-read WGS depth of coverage), and LOH endpoint by short-read WGS were carried out following procedures described previously (Zhang et al. 2013; Heasley et al. 2021). Summaries of results from these genomic analyses are provided in Tables S1 and S3. Multiplex PCR was carried to identify the translocation partners in the *URA-RA3* NAHR competition assays using the Qiagen Multiplex PCR plus kit (Cat. 206152) using the oligonucleotide primers described in Table S5. All WGS dataset generated for this study are deposited and publicly available through in Sequence Read Archive under ascension number PRJNA941438.

## Supporting information

AllSupplementalTables

## ACKNOWLEDGMENTS

Genome stability research in the JL Argueso laboratory was broadly supported by the National Institutes of Health award R35GM11978801. SAM and MJH were supported as fellows through the Predoctoral Training in Quantitative Cell & Molecular Biology from the National Institutes of Health award T32GM132057.

## AUTHOR CONTRIBUTIONS

All authors contributed to the experimental work and data collection. SAM, MJC, JAS, CDS, RSS, MJH, CMP, DAC, RAW and JLA contributed to the experimental design and data analyses. SAM and JLA wrote the manuscript.

